# Dissecting the contribution of recent reward versus recent performance history on cognitive effort allocation

**DOI:** 10.1101/2025.07.08.663681

**Authors:** F.S. Spronkers, R.S. Koolschijn, N.D. Daw, A.R. Otto, H. E.M. den Ouden

## Abstract

An extensive body of literature has shown that humans tend to avoid expending cognitive effort, just like for physical effort or financial resources. How then, do we decide whether to put this effort in? Decision-making not only involves choosing our actions, but also the meta-decision of how much cognitive effort to invest in making this choice, weighing the costs of cognitive effort against potential rewards. Popular recent theories, grounded in the field of reinforcement learning, suggest that this cost-benefit trade-off can be informed by the opportunity costs of effort investment, which the brain may approximate by the estimated average reward rate per unit time. It follows from intuition that in a low reward environment, investing cognitive resources in the task at hand will less likely lead to missed opportunities. Recent studies provided support for this idea, showing that people exert more cognitive effort when reward rate is low. Here, we replicate one of the key previous findings but provide an important nuance to this result. Cognitive effort allocation was better explained by participants’ recent performance history (i.e. accuracy rate) than average reward rate. In combination with the observation that participants were insensitive to the reward currently at stake, this invites a reinterpretation of these previous findings and suggests the need for further studies to assess whether environmental richness may indeed serve as a heuristic to modulate cognitive effort allocation.

## Introduction

Typically, our decisions and actions involve an evaluation of required effort and potential reward (Shenhav et al., 2013). Whether we successfully obtain a reward depends not only on our *ability*, but crucially also on our *motivation* to perform the task, guiding how much effort we are willing to invest (Boureau et al., 2015; Kool & Botvinick, 2018; Shenhav et al., 2013). Given that our cognitive resources are inherently limited (Navon & Gopher, 1979), an important question in recent literature has focused on how we allocate cognitive resources (Devine et al., 2021; Otto & Daw, 2019; Westbrook et al., 2020). An influential recent idea is that when deciding on how much cognitive effort to put in, we weigh the potential benefits of expending effort against the associated costs (Boureau et al., 2015; Kool & Botvinick, 2018). However, this immediately raises the question of how these perceived costs and benefits are estimated neurally in a computationally tractable way (Székely & Michael, 2021). Whilst benefits can be straightforwardly operationalised as expected reward magnitude, the metric of ‘costs’ is more complicated. A recent proposal is that these internally estimated costs should include ‘opportunity costs’ – the idea that when we choose to invest our time and effort in one activity, we inherently forego the opportunity to allocate those resources to something else (Kurzban et al., 2013). In this study, we aim to replicate and nuance one of the first empirical studies from an increasing body of research assessing the idea that estimated opportunity costs inform cognitive effort allocation (Otto & Daw, 2019).

Opportunity cost can be approximated by tracking the rewards obtained over time, i.e. the reward rate of the environment (Niv et al., 2007). This is most intuitive in the domain of physical vigour, where faster responses are more costly, but also allow for a larger number of rewards to be obtained. When reward rate is high, the opportunity cost of ‘sloth’ (i.e. of spending time) is particularly high. Supporting this idea, increased reward rate has been associated with speeding of responses, suggested to reflect an increased willingness to exert physical effort to reduce opportunities lost due to spending too much time on a single response (Beierholm et al., 2013; Guitart-Masip et al., 2011).

Like physical effort, we also seek to avoid cognitive effort, as a finite resource where prolonged effort leads to fatigue (Botvinick & Braver, 2015; Navon & Gopher, 1979; Pessiglione et al., 2025). The expected value of control (EVC) theory posits that an individual’s decision to exert (costly) control is based on a cost-benefit analysis aimed to maximize reward while minimizing intrinsically costly effort exertion (Shenhav et al., 2013). Like for physical effort, environmental reward rate many serve as a heuristic proxy to estimate how much cognitive effort is worth investing in a particular task. Indeed, environmental reward rate has been shown to not only impact physical vigour but also cognitive effort exertion across various tasks, ranging from perceptual decision-making to classic cognitive control tasks. Here, when the reward rate was high, subjects became faster and more error-prone, which was interpreted as a withdrawal of cognitive effort. Thus, expenditure of cognitive effort appeared to be modulated by the opportunity cost of time (Devine et al., 2021; Lin et al., 2022; Otto & Daw, 2019).

With growing interest in the idea that environmental reward rate may serve as a computational proxy to opportunity costs, and can thus be used to modulate cognitive effort allocation (Devine et al., 2021; Lin et al., 2022; Niv et al., 2007; Otto & Daw, 2019), this study aims to assess the robustness of these observations, specifically replicating Otto & Daw (2019). This study combined the Simon task, a simple, well-established cognitive control task (Simon & Berbaum, 1990) with a reward incentive manipulation. We replicate the previous finding that participants increased cognitive control under low compared to high reward rate (Otto & Daw, 2019), quantified as increased accuracy while leaving response speed unaltered. Crucially, however, we find that accuracy rate (i.e. the long-running average accuracy) better predicted performance than average reward rate. Combined with the observation that speed and performance were insensitive to the reward at stake on the current trial, this invites a reinterpretation of previously identified reward rate effects. We discuss important implications of these findings and outline future directions for investigating cognitive effort allocation.

## Materials & Methods

This study was pre-registered on OSF under https://doi.org/10.17605/OSF.IO/QET4M. The first objective of the current study was to replicate the results from Otto & Daw (2019). Subsequently, we conducted a set of analyses to further characterise the replicated average reward rate effect.

### Participants

Sixty-nine healthy volunteers (mean age = 23, SD = 3.8, range = 18-35, 47 women, 61 right-handed) participated in the study, which was approved by the local ethics committee (ECSW2017-2306-520). Participants gave written informed consent prior to beginning the task.

They received a monetary reward of €7.50 plus a performance-based bonus of up to €2.50 (mean €1.96, SD €0.24). We excluded one participant from data analysis because they missed more than 25 percent of trials during the main phase of the experiment, leaving 68 participants for the analysis.

### Simon task

Participants performed a Simon task with reward magnitude manipulation(Otto & Daw, 2019). The task was presented using PsychoPy 3.2.4 for Python 3.7.5 (Peirce et al., 2019) on a BenQ XL2420Z computer screen with a 1920×1080 resolution and a 60 Hz refresh rate. On each trial, participants saw a blue or green circle on either the left or right side of the screen (Fig 1A). The stimulus colour indicated whether participants should press the left or right button on a button box. In 75% of trials (‘congruent’ trials), the stimulus location matched the lateralization (left/right button) of the correct hand response, while in 25% of trials (‘incongruent’ trials), the stimulus appeared on the opposite side of the correct hand response. Before each stimulus, the reward that could be gained for correct responses, ranging between 1 and 100 points, was shown for 750-1250 ms (the interstimulus interval). Participants were instructed to respond before the coloured stimulus disappeared. After the stimulus feedback interval (500ms), feedback was presented for 1000 ms. indicating whether the participant was correct or not or responded too late. Only correct, on-time responses were rewarded (Fig 1B). For one participant, one trial was excluded because the response time was more than 3 standard deviations away from the mean. Relative to the stimulus presentation in Otto & Daw (2019), we shortened the presentation time by 100 ms. because participants performed at ceiling level in pilots using the original timings (accuracy congruent trials: 95.0% SD= 3.1, range: 90.4% - 99.5%, accuracy incongruent trials: 87.8% SD = 9.3, range: 63.0% - 99.1%, see Supplemental Materials for analysis of the pilot dataset). In 90% of trials, the stimulus was presented for 500 ms. In the remaining 10% of trials, the stimulus was presented for 400 ms. to ensure that participants continued to pay attention and respond as quickly as possible.

**Figure 1.**
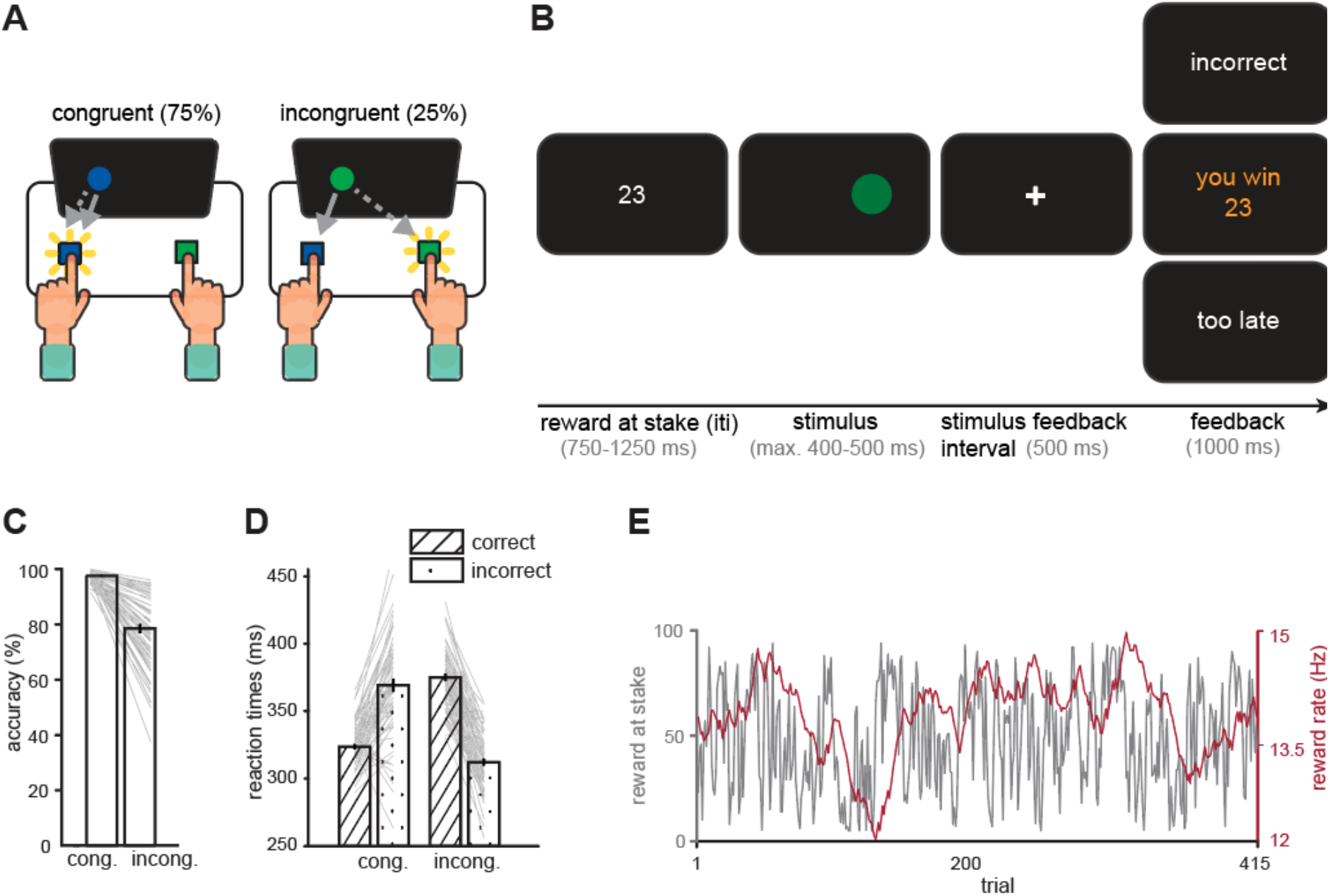
Task design. **A**. Simon task. Participants were required to respond according to the colour of the stimulus on the screen. On congruent (cong.) trials, the coloured stimulus appeared on the side of the required hand response, while on incongruent (incong.) trials the stimulus appeared on the opposite side of the hand response. **B**. Participants were presented with the reward at stake (iti, 750-1250 ms.), after which the stimulus appeared, and they had to respond within 400 ms. (10% of trials) or 500 ms. (90% of trials). After the stimulus feedback interval, participants received feedback. For correct responses this was the number of points presented at the start of the trial, and for incorrect or too late trials they received 0 points. **C**. Accuracy for congruent and incongruent trials, error bars represent standard error of the mean. Participants are more accurate on congruent trials than incongruent trials. **D**. Reaction times for correct congruent, incorrect congruent, correct incongruent and incorrect incongruent trials. Participants are faster on congruent trials when they make a correct response, but faster on incongruent trials when they make a mistake. **E**. Example reward sequence and the corresponding reward rate sequence. The reward rate was calculated from the rewards received, with a learning rate of 0.0031 (Otto & Daw, 2019).

Importantly, to manipulate the reward rate during the experiment, the amount of reward that could be won for a correct response was varied according to three predetermined pseudo-random sequences, previously used in Otto & Daw (2019; Fig S1). The sequences were counterbalanced across participants. These sequences were constructed such that participants could earn roughly the same amount of reward on average over 500 trials (average reward magnitude each reward sequence was [50.4; 48.9; 47.3] points).

Participants first practiced 10 trials, after which they completed 120 preliminary trials without reward at stake. They were then instructed about the rewards they could earn for correct responses. Participants performed the task with the stake manipulation for 20 minutes, completing as many trials as possible. At the end of the task, they received a monetary performance bonus based on their accumulated score.

### Replication analysis

All analyses were performed in R v4.1.0 using Rstudio v1.4.1717. We performed mixed-effects linear regressions and logistic regressions using the package lme4 (Bates et al., 2015). P-values were calculated using the function “mixed” in the afex package (Singmann et al., 2024), except for the sensitivity analyses (see below) where likelihood ratio tests were used from the lme4 package due to the long computation time required by the “mixed” function. Figures were made with Matlab R2024a.

### Data inclusion

Replicating Otto & Daw (2019) we removed the following trials: (i) all trials where participants missed the response deadline; (ii) the first 10 trials for each participant as a ‘burn-in’ period to allow participants to form an estimate of the reward rate; and (iii) trials with reaction times more than three standard deviations away from the participant’s mean reaction time, as these likely reflect lapses of attention.

### Average reward rate calculation

To calculate the reward rate per second (*ARR*), we used the update rule following (Constantino & Daw, 2015):

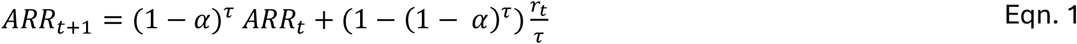

Here, *r*_*t*_ is the received reward, τ is the time elapsed since the last update, and α is the learning rate per second.

For an exact replication of experiment two in Otto & Daw (2019), we used a learning rate of 0.0031 for all participants (which there was estimated on data from a different task paradigm). In addition, following the method later used by Lin et al. (2022) using the same task, we estimated the learning rate on response time (RT) data of correct, congruent trials. The rationale for this approach is to independently estimate the reward rate learning rate on RT data and subsequently assess the effect of reward rate in the choice data. Here, we assume that reward rate modulation of previously observed invigoration of physical effort (Guitart-Masip et al., 2011; Otto & Daw, 2019) would also shape cognitive effort. An example reward rate sequence is shown in Figure 1E.

We estimated a single learning rate α using a general-purpose Nelder-Mead optimization in R, using the following model:

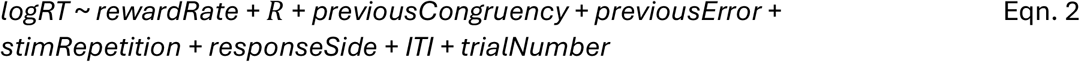

Extra factors in addition to reward rate were included to capture as much variance as possible, thereby eliminating the influence of these nuisance variables (Beierholm et al., 2013; Otto & Daw, 2019). All binary variables were factorized, and continuous variables were z-scored. RTs were log-transformed. *R* is the reward at stake; *previousError* and *previousCongruency* represented events on the previous trial, to account for adaptations like post-error slowing and pro-active control allocation (Ridderinkhof, 2002); *StimRepetition* is whether the same response was required on the previous trial, where stimulus repetition is expected to lead to a faster response; *ResponseSide* represented which button the participant was supposed to press to capture possible response biases; *trialNumber* was included to capture speeding or slowing over time due to learning or fatigue respectively. The intertrial interval (*ITI*) is part of the reaction time model because a longer ITI and therefore longer preparation time could speed up responding. Note that during the ITI, the reward at stake is presented. Fitting this regression model while optimizing for the learning rate resulted in a best-fitting learning rate of 0.0042. The regressions described below were performed with both the fixed (0.0031; Otto & Daw, 2019) and fitted (0.0042) learning rates. Note that below, we further explore the robustness of the results under learning rates.

### Regression analyses

To assess whether reward rate affects accuracy, where we hypothesized that increased reward rate would lead to decreased performance, especially on incongruent trials, we conducted a mixed-effects regression using the following model:

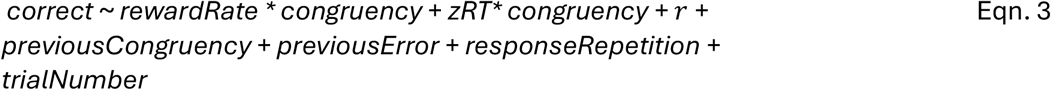

Previous congruency, previous error and response repetition were included in the models to account for potential performance adaptation as a function of the previous trial. Specifically, we expected participants to make fewer mistakes if the previous trial was incongruent, if they made a mistake on the previous trial, or if the stimulus was repeated and thus the current trial required the same response as the previous trial. Lastly, trial number was included to capture potential (linear) task learning or alternatively fatigue effects over the course of the task.

We adapted the model used by Otto & Daw (2019) to include the interaction between the reward rate and congruency because cognitive control/ mental effort is only required on the incongruent trials, and thus we expected reward rate to primarily affect those incongruent trials, as also reported in previous studies (Lin et al., 2022; Mittelstädt et al., 2023; Otto & Daw, 2019). To confirm the specificity of the effect to the incongruent trials, we analysed congruent and incongruent trials separately as a post-hoc follow-up analysis. Our key hypothesis is that under high reward rates, cognitive effort decreases. This predicts not merely speeding and concomitant reduction in accuracy (i.e. a shift along the speed-accuracy trade-off function - SATF), but rather a downward shift of the SATF, i.e. reduced performance given the same RT. To account for potential SATF effects, we included Z-scored RTs as a dependent variable. This enables us to isolate effects of reward rate on accuracy over and above a shift along the SATF as modulation of cognitive effort.

We furthermore repeated the RT analysis in a mixed linear regression model now including RTs for correct responses on both congruent and incongruent trials, to assess whether there are indeed effects of reward rate on physical vigor (i.e. RTs). We used the same model as Otto & Daw (2019):

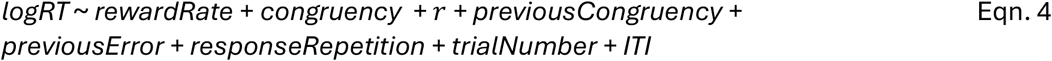

We expected to reward rate to increase response vigor across congruent and incongruent trials. Following the findings of Otto & Daw (2019), we expected no effects of reward at stake. We further expected to see slower RTs when the previous trial was congruent (Gratton et al., 1992), post-error slowing, faster responding for congruent than incongruent trials, and speeding if the current trial required the same response as the previous trial. Finally, we predicted shorter RTs with increased trial number (task learning) and ITIs (readiness).

### Exploratory analyses – Learning rate sensitivity analysis

It has been shown previously that for relatively simple learning models, the exact value of a learning rate may affect the amplitude but not the shape of the learning curve, and thus not affect results of regression analyses using the learning curve as a covariate (Wilson & Niv, 2015). To quantify this, we assessed the degree of co-variance of reward rates at different learning rates, by conducting a grid search over reward rate learning rates in the range of [0 1]. Here, we oversampled the lower learning rates, as previously observed effects of reward rate were in this regime (Beierholm et al., 2013; Guitart-Masip et al., 2011; Otto & Daw, 2019). We increased step size with increasing learning rate values, taking 100 evenly spaced samples for each of the following intervals: [1 *10^−5^ - 1 *10^−4^],[1 *10^−4^ - 1 *10^−3^], [1 *10^−3^ - 1 *10^−2^], [0.01 – 0.1], [0.1 -1].

To also examine whether the reward rate effect depended on the specific learning rate, we computed the reward rate effect across a range of learning rates, focusing on incongruent trials (Eqn. 3), which were sensitive to reward rate.

### Exploratory analyses – Characterizing reward rate effects

We conducted several exploratory analyses to further characterize the replicated effect of reward rate on cognitive effort. First, we examined in more detail the finding that there was no significant effect of reward at stake on task performance (both RT and accuracy), replicating previous studies (Beierholm et al., 2013; Guitart-Masip et al., 2011; Lin et al., 2022; Otto & Daw, 2019). This is puzzling given the extensive body of literature showing that stakes can influence task performance in a rewarded Stroop task (Aarts et al., 2014; Schmidt et al., 2012), although this may only hold for longer stake horizons (Kukkonen et al., 2025).To relate long-running reward rates to instantaneous reward-on-offer effects, we therefore first examined whether the last received reward also influenced performance. We replaced the reward rate variable in the accuracy regression (Eqn. 3) with the reward received at the previous trial, analysing the incongruent trials only, as only these trials showed an effect of reward rate.

Given the absence of an immediate effect of the reward at stake, we assessed whether participants were sensitive at all to the long-running average of the magnitude of received rewards, or that behaviour rather tracks whether they receive rewards or not (in this case perfectly correlated with response accuracy). Therefore, we calculated the “accuracy rate” using the same updating rule as in Eqn. 1, but with binary coding of the received reward, using the same learning rate of 0.0031 (but also repeating the grid search across learning rates, as described below). We reran the accuracy- (Eqn. 3) and RT-predicting (Eqn. 4) regressions with this accuracy rate.

To assess whether accuracy rate effects were driven by external rewards or also generalize to a non-rewarded task setting, we tested if the accuracy rate affected performance during the 120 preliminary trials, where no rewards were presented but participants received accuracy feedback. Finally, we assessed whether reward rate explained variance over and above the accuracy rate, which dissects out the *magnitude* aspect of received rewards. To this end, we repeated the regression analyses in Eqn. 3 and Eqn. 4, including both the accuracy rate and orthogonalized reward rate in the model.

## Results

### Data inclusion

Participants performed 421 trials on average (SD 0.3, range 414 - 428) of the Simon task during 20 minutes of allotted time. Participants missed 23 trials on average (SD 1.6, range 5-65) during the main phase of the experiment. On average 4.1% of 500 ms. deadline trials were missed, and 15.5% of 400 ms. deadline trials were missed. Note that any responses after the deadline were not recorded and thus not analysed.

### Replication of previously reported reward rate effects on cognitive effort exertion

The aim of this study was to replicate the finding that participants exert more cognitive effort when reward rate is low compared to when the reward rate is high (Devine et al., 2021; Lin et al., 2022; Otto & Daw, 2019). Note that throughout this section, we report statistics from two mixed-effect regression analyses, both using the learning rate reported in Otto and Daw (2019) (0.0031, α1) and using the learning rate fitted to RTs on correct, congruent trials the present dataset (0.0042, α2, see *Methods*). In line with previous studies using the Simon task, participants made more mistakes (Figure 1C, β_α1_ = 1.5, β_α2_ = 1.5, p_α1_ < 2×10^−16^, p_α2_ < 2×10^−16^) and were slower (Figure 1D, β_α1_ = -25.2, β_α2_ = -25.2, p_α1_ < 2×10^−16^, p_α2_ = 2×10^−16^) on incongruent trials. Furthermore, participants were less accurate (Fig S2A, *previousError:* β_α1_ = -0.4, β_α2_ = -0.4, p_α1_ = 7×10^−13^, p_α2_ = 1×10^−14^) and slower (Fig S2B, *previousError:* β_α1_ = 8.7, β_α2_ = 8.7, p_α1_ = 1×10^−12^, p_α2_ = 1×10^−12^) after making a mistake. Lastly, they were less accurate (Fig S2C, *previousCongruency:* β_α1_ = -0.3, β_α2_ = -0.3, p_α1_ = 2×10^−7^, p_α2_ = 2×10^−7^) and faster (Fig S2D, *previousCongruency:* β_α1_ = -4.7, β_α2_ = -4.7, p_α1_ < 2×10^−16^, p_α2_ = 1×10^−13^) following an incongruent trial.

Reward rate modulated accuracy as a function of congruency (Figure 2A-C, *rewardRate*congruency*: β_α1_ = 0.09, β_α2_ = 0.09, p_α1_ = 0.03, p_α2_ = 0.03, Table 1, 2 and S1). Post-hoc simple tests showed that on incongruent trials, but not congruent trials, people made more mistakes when reward rate was high (*rewardRate* - congruent trials: β_α1_ = -0.005, β_α2_ = -0.0003, p_α1_ = 0.9, p_α2_ = 0.99; incongruent trials: β_α1_ = -0.1, β_α2_ = -0.1, p_α1_ = 0.006, p_α2_ = 0.007). Unlike in Otto & Daw (2019), there was no effect of reward rate on reaction times (Fig 2D+E and Table 3, *rewardRate:* β_α1_ = -0.05, β_α2_ = -0.1, p_α1_ = 0.9, p_α2_ = 0.9, see also Figure S3 further exploring the absence of an effect of reward rate on RT). These results indicate that reward rate did not shift performance along a speed-accuracy trade-off (e.g. increasing speed and simultaneously decreasing accuracy) but rather shifted the speed-accuracy trade-off function itself, where under a higher reward rate, for the same RT participants would make more mistakes (Fig 2F). Consistent with previous studies (Beierholm et al., 2013; Devine et al., 2021; Guitart-Masip et al., 2011; Lin et al., 2022; Otto & Daw, 2019), there was no significant effect of reward at stake on accuracy or RT (Table 1&3).

**Table 1.**
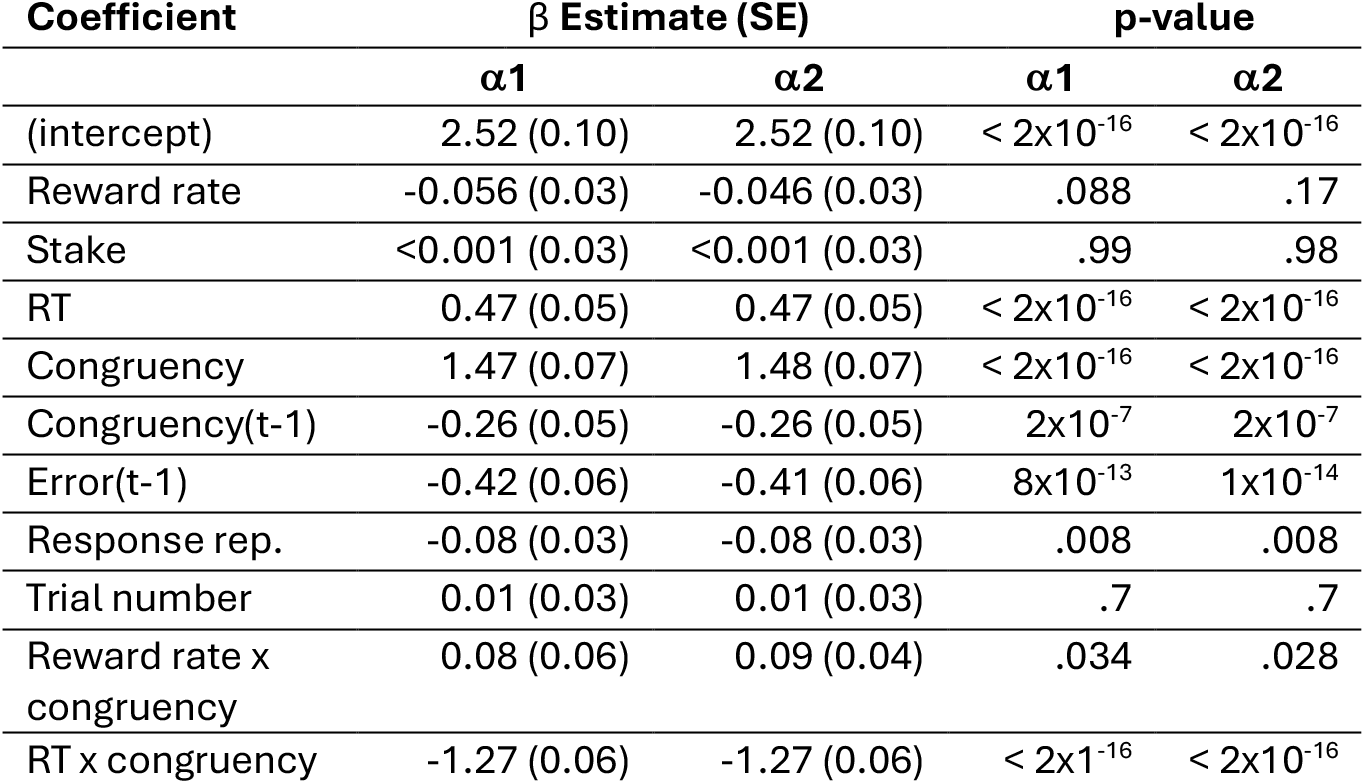
Accuracy. Results from the mixed-effects logistic regression analysis examining effects on accuracy for all trials. Results are consistent across two learning rates: α1 = 0.0031, α2 = 0.0042.

**Table 2.** Accuracy - incongruent trials. Results from the mixed-effects logistic regression analysis examining effects on accuracy for the incongruent trials. Results are consistent across two learning rates: α1 = 0.0031, α2 = 0.0042.

| Coefficient | $\beta$ Estimate (SE) | | p-value | |
| --- | --- | --- | --- | --- |
| | $\alpha 1$ | $\alpha 2$ | $\alpha 1$ | $\alpha 2$ |
| (intercept) | 2.13 (0.17) | 2.12 (0.17) | $< 2 \times 10^{-16}$ | $< 2 \times 10^{-16}$ |
| Reward rate | -0.12 (0.04) | -0.11 (0.04) | .006 | .007 |
| Stake | 0.008 (0.04) | 0.01 (0.04) | .9 | .8 |
| RT | 1.86 (0.11) | 1.86 (0.11) | $< 2 \times 10^{-16}$ | $< 2 \times 10^{-16}$ |
| Congruency(t-1) | -0.86 (0.07) | -0.86 (0.07) | $< 2 \times 10^{-16}$ | $< 2 \times 10^{-16}$ |
| Error(t-1) | -0.70 (0.11) | -0.70 (0.11) | $2 \times 10^{-10}$ | $5 \times 10^{-11}$ |
| Response rep. | -0.10 (0.04) | -0.09 (0.04) | .03 | .02 |
| Trial number | 0.08 (0.05) | 0.09 (0.05) | .07 | .06 |

**Table 3.** Reaction times. Results from the mixed-effects linear regression analysis examining effects on reaction times for all correct trials. Results are consistent across two learning rates: α1 = 0.0031, α2 = 0.0042.

| Coefficient | $\beta$ Estimate (SE) | | p-value | |
| --- | --- | --- | --- | --- |
| | $\alpha 1$ | $\alpha 2$ | $\alpha 1$ | $\alpha 2$ |
| (intercept) | 350.4 (3.3) | 350.4 (3.3) | $< 2 \times 10^{-16}$ | $< 2 \times 10^{-16}$ |
| Reward rate | -0.30 (0.58) | -0.35 (0.59) | .6 | .6 |
| Reward at stake | -0.46 (0.35) | -0.45 (0.35) | .2 | .2 |
| Congruency | -17.72 (0.70) | -17.71 (0.69) | $< 2 \times 10^{-16}$ | $< 2 \times 10^{-16}$ |
| Congruency(t-1) | -4.69 (0.51) | -4.69 (0.51) | $< 2 \times 10^{-16}$ | $1 \times 10^{-13}$ |
| Error(t-1) | 8.72 (0.98) | 8.71 (0.98) | $1 \times 10^{-12}$ | $1 \times 10^{-12}$ |
| Response rep. | 1.63 (0.69) | 1.63 (0.69) | .02 | .02 |
| Trial number | 0.59 (0.67) | 0.60 (0.67) | .4 | .4 |
| ITI | -5.56 (0.48) | -5.56 (0.48) | $< 2 \times 10^{-16}$ | $< 2 \times 10^{-16}$ |

In summary, the analyses above show nearly identical findings for both the learning rate used by Otto & Daw (2019) and the learning rate estimated on correct, congruent trials in the current dataset. This suggests that the precise learning rate may not be critical to characterising the effects of reward rate. To examine if the effect of reward rate on behaviour depends on the exact learning rate, we conducted a systematic sensitivity analysis, where we estimated reward rate effects on accuracy for a range of learning rates (range 0 to 1), for incongruent trials only (see *Methods*). Reward rate significantly predicted accuracy for learning rates below 0.007 (Fig 3A), which showed high cross-correlation (Fig 3B). Thus, in the learning rate domain where reward rate reflects a long-range average, reward rate significantly explained cognitive effort. In line with the observation that long-range integration of reward history affected accuracy, there was no significant effect of reward received on the preceding trial on accuracy (β = 0.07, p=0.14, Fig 3C).

**Figure 3.**
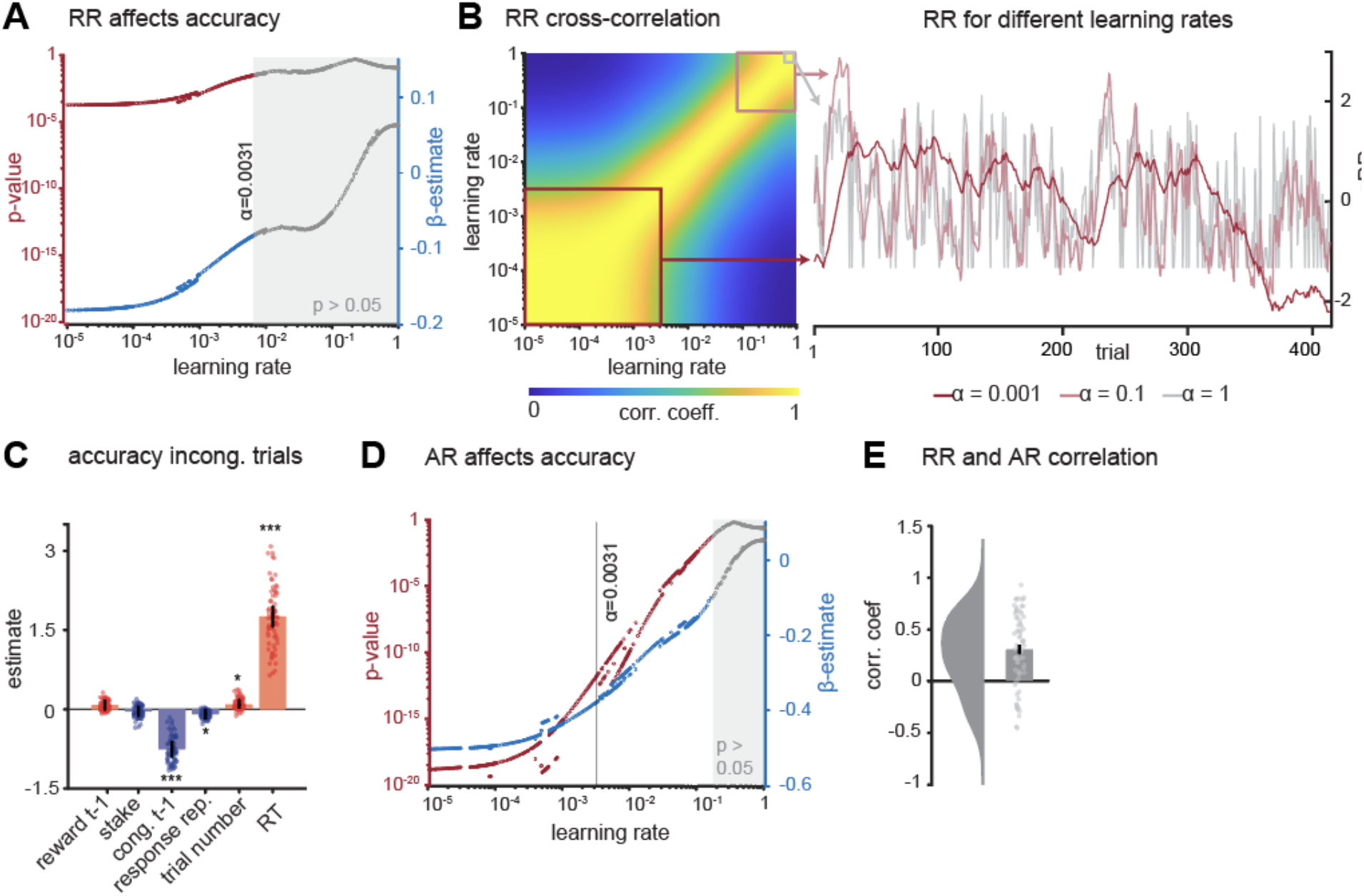
Exploring the effect of learning rate on reward rate (RR) and accuracy rate (AR) effects. **A**. Effect of reward rate on incongruent trials, with p-values (red) and estimates (blue) for reward rate computed for a range of learning rates between 0 and 1. The effect of reward was significant for learning rates below 0.007. P-values and estimates are shown in grey where the effect of reward rate was p>0.05. **B**. Cross-correlation of reward rate dynamics as a function of learning rate. Reward rate sequences with a learning rate below 0.007 significantly explained cognitive effort exertion and are highly intercorrelated (red square). Reward rates in the range of 0.2-1 are also highly intercorrelated, indicating that a high learning rate results in a reward rate that is effectively reflecting the reward received on the previous trial (pink square). On the left is an example of corresponding reward rate sequences calculated with three different learning rates: 0.001 (red), 0.1 (pink) and 1 (grey). **C**. Regression coefficients of a mixed-effects logistic regression analysis on accuracy with the reward received on the previous trial, which represents a learning rate of 1. There was no significant effect of reward received on the previous trial on accuracy. **D**. Effect of accuracy rate (AR) on incongruent trials, for learning rates ranging 0 to 1 to compute AR. The effect of accuracy rate significantly explains accuracy for learning rates < 0.16. P-values and estimates are shown in grey when the effect of AR is not significant. **E**. Correlation of reward rate and accuracy rate, computed at a learning rate of 0.0031. While on average RR and AR are positively correlated, in some participants they are negatively correlated. These participants thus make fewer mistakes in a low reward environment. *** p<0.001; ** p<0.01; * p<0.05.

### Past performance predicts behaviour better than reward history

While reward rate predicted cognitive effort, stake magnitude on the current trial did not (Figure 2C, Table 2). As reward rate integrates rewards obtained for correct responses, and this lack of a stake effect could suggest that participants did not process reward magnitude, we assessed whether omitting reward magnitude information from the reward rate calculation affected the reward rate effect. In other words, we examined the extent to which the effect of reward rate might be driven purely by whether participants received a reward, but not how large. As rewards were received only on correct trials, this then boils down to the effect of recent history of performance. Participants made significantly fewer mistakes when the accuracy rate was lower (Fig 4A-B, Table S2, *accuracyRate:* β =-0.4, p <2×10^−16^), while there was no significant effect of accuracy rate on RT (Fig 4D+E, Table S5, accuracyRate: β =-0.3, p =0.6). While there was a trend for an interaction between accuracy rate and congruency (accuracyRate*congruency: β =0.06, p =0.09), the effect of accuracy rate was significant for both congruent and incongruent trials (Table S3+S4, accuracyRate - congruent: β =-0.3, p = 1.0×10^−6^; incongruent: β =-0.4, p < 2×10^−16^, Fig 4C). Similar to reward rate, the effect of accuracy rate on accuracy was significant for a range of low learning rates (Fig 3D). Taken together, an increase in accuracy rate was associated with a downward shift of the speed-accuracy trade-off (Fig 4F), that was much stronger (in terms of effect size and significance) than the effect of reward rate. Indeed, models with accuracy rate outperformed models with reward rate for both accuracy and RT models in terms of estimated model evidence (Fig 5).

**Figure 4.**
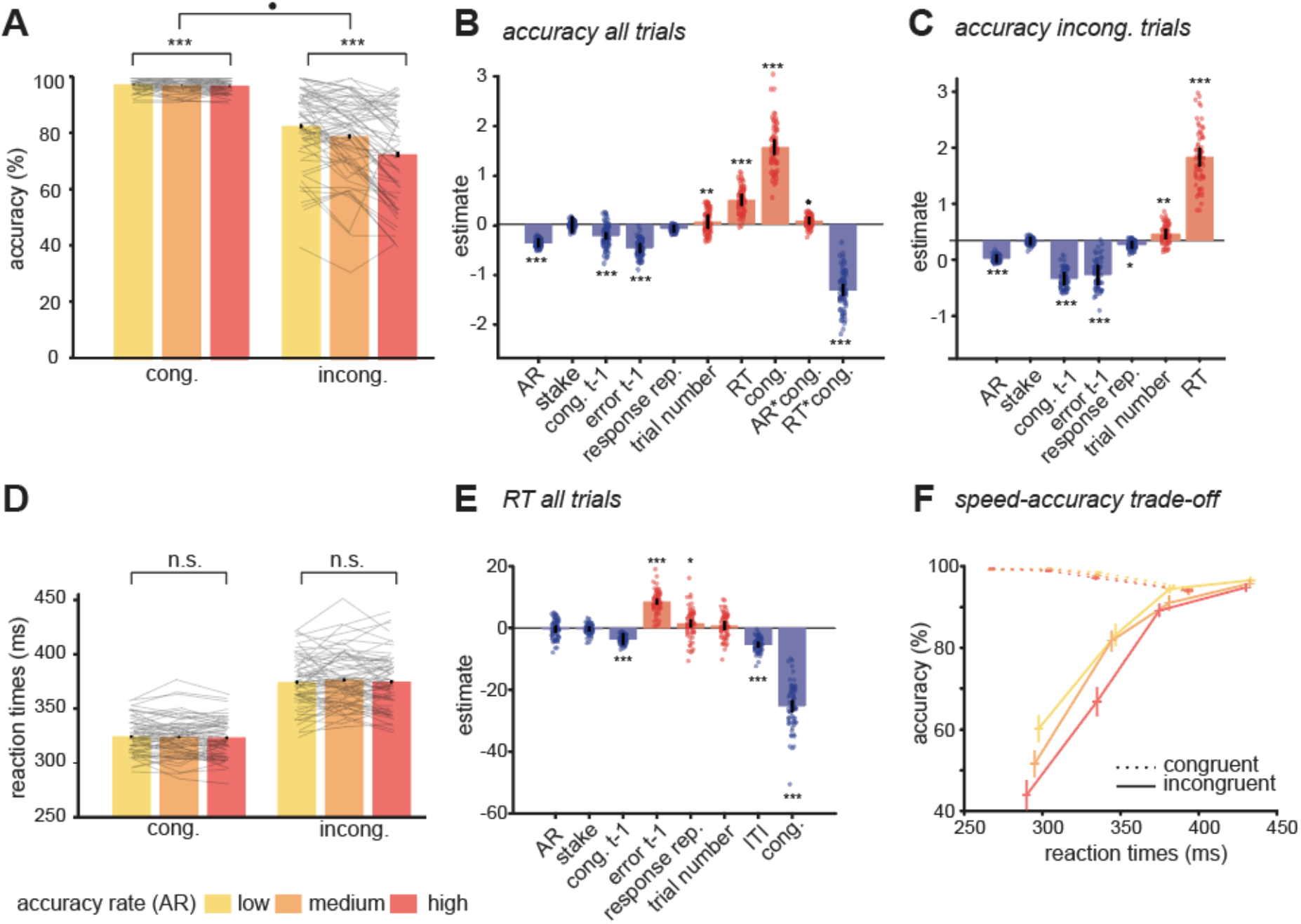
Accuracy rate affects task performance. **A**. Accuracy on congruent and incongruent trials as a function of reward rate (tertile split for illustration only). On incongruent trials, participants were significantly more accurate when the accuracy rate (AR) was low compared to when the accuracy rate was high. On congruent trials, performance was nearly at ceiling. **B-C**. Regression coefficients from the accuracy mixed-effects logistic regression for all trials (B) and incongruent trials only (C). Importantly, there was no effect of reward at stake on the current trial. The accuracy rate affected performance on all trials: participants were more accurate when the accuracy rate was low. **D**. RTs on congruent and incongruent trials as a function of accuracy rate. Accuracy rate did not significantly affect RT. **E**. Regression coefficients from the RT mixed-effects linear regression for all trials. RTs were faster for congruent trials, when the previous trial was congruent (cong. t-1), and when inter-trial interval (ITI) was longer. RTs increased when participants had made an error on the previous trial and when the same button was pressed (response rep.). There was no effect of accuracy rate or reward at stake on RT. **F**. There was a downward shift of the speed-accuracy trade-off with increasing accuracy rate for the incongruent trials only. *** p<0.001; ** p<0.01; * p<0.05.

**Figure 5.**
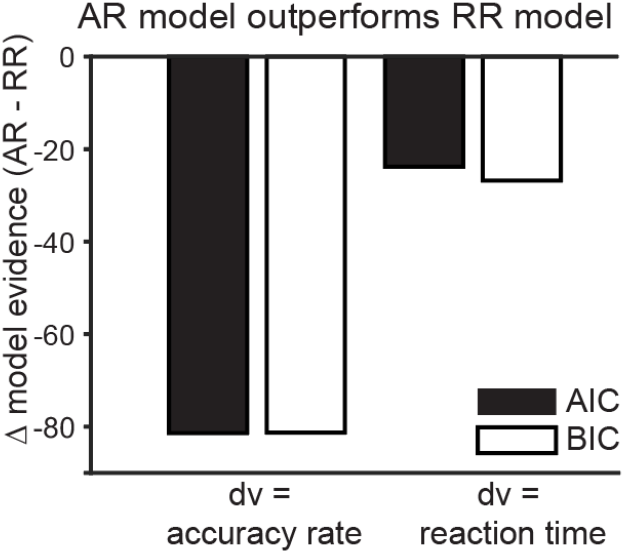
Accuracy rate better explains behaviour than reward rate. Difference in model evidence between the regression with the accuracy rate minus the model with reward rate regressor. We used the learning rate of 0.0031 (Otto & Daw, 2019) to calculate the accuracy and reward rates. Left: model with accuracy as dependent variable. Right: model with reaction time as dependent variable. For both the accuracy and reaction time regression analyses, the accuracy rate model outperforms the reward rate model, evident from negative AIC and BIC delta values.

Given that the accuracy rate regressor and reward rate regressors differ only in terms of ignoring or including reward magnitude information, we first quantified the degree of correlation of these regressors, and next whether addition of reward magnitude information explains additional, unique variance. While reward rate on average correlated positively to reward, the strength of this correlation varied highly between participants, explaining on average 9% of variance, and was even negative for some participants (Fig 3E, mean R=0.29, SD=0.041; range [-0.44–0.88]). This relatively low correlation is a result of the fluctuating reward magnitude manipulation and confirms that effects of reward and accuracy rate can in principle be identified separately. However, when orthogonalising accuracy rate and reward rate, reward did not explain additional variance over and above the effect of accuracy rate (Table S6).

There are at least two explanations for the finding that behaviour is better explained by accuracy than reward rate. First, participants may track the presence of rewards without considering their magnitude. Alternatively, they might track task performance rather than the rewards themselves. These two options can be dissociated by examining whether the effect of the accuracy rate persists when participants do not receive reward. We therefore conducted the same regression analyses on the 120 preliminary trials where no stakes or rewards were presented to participants (only correct / incorrect feedback, Table S7-S10). Even when no reward was presented, participants made significantly fewer mistakes when the accuracy rate was lower (accuracyRate: all trials: β =-2.3, p = 6×10^−7^; congruent: β =-2.5, p =0.002 incongruent: β =-2.4, p =0.009), showing that participants track performance rather than binary reward. Also, for RT, accuracy rate effects on preliminary trials mirrored the main findings that participants were faster when accuracy rate was high (accuracyRate: β =-17.9, p =0.002).

Taken together, our findings replicate those of Otto & Daw (2019), showing that participants exert more cognitive effort when the reward rate is lower. While we did not replicate the previously observed slowing of responses for lower reward rates (Otto & Daw 2019), this absence of a slowing effect aligns with findings from a later study (Lin et al., 2022). Crucially, however, we demonstrate that the rate of correct responding predicts performance better than a reward rate integrating the magnitude of the received reward.

## Discussion

Decision-making involves not only selecting an action but also determining how much of our limited cognitive resources to allocate to this decision process. To make this ‘meta’-decision, we must weigh the costs of cognitive effort against potential rewards. In the present study, we aimed to replicate the findings of a set of cognitive control experiments (Otto & Daw, 2019, see also Devine et al., 2021, and Lin et al., 2022) testing the idea that we track the average reward rate of the environment as a proxy for the opportunity cost of time (Niv et al., 2007), in turn directing cognitive effort allocation decisions. Replicating these previous findings, we found that participants exerted more cognitive effort when (average) environmental reward rate was low, and withheld effort when the environmental reward rate was high. Crucially, however, this modulation of cognitive effort did not depend on reward magnitude; rather, cognitive effort was better explained by the recent history of performance accuracy. Below we discuss both theoretical and empirical implications of these findings.

Previous work proposed interpreting the observed modulation of performance by environmental reward rate as reflecting sensitivity to the fluctuations in opportunity costs of time (Otto & Daw, 2019; Devine et al., 2021; Lin et al., 2022), following similar theoretical reasoning for optimizing physical effort (i.e., vigor; Beierholm et al., 2013; Niv et al., 2007). In short, when the average environmental reward rate is high, opportunity costs of time are similarly high, and thus speed is of the essence. However, in the context of cognitive rather than physical effort, it is less clear whether opportunity cost of time is the primary factor being optimized. Performance on cognitive control tasks, like the Simon task, is well-characterized by a so-called speed-accuracy trade-off function (SATF), where speeding (‘spending less time thinking’) leads to lower accuracy, while slowing increases accuracy. In the original study (Otto & Daw 2019), high-reward rate environments jointly induced both speeding and decreases in accuracy, in line with the interpretation of reducing opportunity costs of time. However, modulations of performance by reward rate could not be explained purely by a shift along the SATF (Otto & Daw 2019), instead showing modulation of accuracy even after accounting for shifting along the SATF.

Furthermore, a later study (Lin et al., 2022) as well as the current study did not replicate reward rate modulating speed, but rather only observed modulations in accuracy. Here, low reward rates are associated with increased accuracy, in the absence of modulating speed, suggestive of an increase in cognitive effort *per unit time*. In other words, investing more cognitive resources could shift the speed-accuracy curve so that one achieves higher accuracy by ‘thinking harder’ for the same amount of time. Thus, an alternative interpretation is that rather than the opportunity costs of time, the average reward rate could modulate opportunity costs of cognitive resource investment. Here, a low reward rate signals the need to secure the limited rewards available, thereby motivating increased effort. Conversely, a high reward rate indicates that the cost of missing an opportunity due to reduced effort investment is minimal, as another high-reward opportunity will be available soon.

To investigate whether the experienced reward richness of the environment modulates cognitive effort, we manipulated the reward rate of the environment by fluctuating the reward at stake on a trial-by-trial basis. Manipulating the reward at stake has been proven effective in inducing fluctuations in performance (increasing accuracy and/or speed) on the current trial across experimental scenarios (e.g., Adcock et al., 2006; Hofmans et al., 2020; Knutson et al., 2003; Krawczyk et al., 2007; Schmidt et al., 2012; see Burton et al., 2021, for a meta-analysis demonstrating effects of stake in inhibitory control tasks). However, in our study, as well as in previous studies employing the same fluctuating reward rate manipulation (Beierholm et al., 2013; Devine et al., 2021; Guitart-Masip et al., 2011; Lin et al., 2022), reward at stake did not affect performance, which suggests that participants failed to process stake information. This, in turn, may underlie our finding that reward history did not explain cognitive effort exertion over and above recent performance history (‘accuracy rate’). Crucially, the accuracy rate effect was also present during the preliminary trials, where no rewards were presented. This absence of a reward rate effect (over and above accuracy rate) merits a closer look at the interpretation that the reward (or performance) rate modulation is driven by minimizing opportunity costs of cognitive effort.

The finding that reward rate had no explanatory power over and above recent performance history, combined with the observation that performance was not sensitive to the reward at stake on the current trial, suggests that a factor other than environmental richness explains variability in cognitive effort in the current paradigm. One possibility is that the accuracy rate metric reflects integration of binarized rewards, omitting reward magnitude information, which may not have been salient enough for participants to utilize in effort (see also supplemental materials). However, the presence of this accuracy rate effect in preliminary trials (where no reward feedback was provided) suggests that tracking of errors, rather than rewards, is the most likely candidate to modulate cognitive effort. Thus, rather than reflecting a proxy of opportunity costs of cognitive effort, this accuracy rate might function as a performance-monitoring signal: participants might dynamically adjust cognitive effort to keep within a reasonable error range, signalling the need to increase effort when dropping too low, whereas a high accuracy rate might signal that cognitive resources can be preserved because enough reward is being obtained. A final possibility is that the observed effects are driven by fatigue, as a high accuracy rate would require sustained cognitive effort, leading to more errors following a period of high accuracy. However, as performance improved (rather than degraded) over time, this suggests at the very least that there are no linear effects of fatigue decreasing performance.

In conclusion, while we replicate previous findings (Devine et al., 2021; Lin et al., 2022; Otto & Daw, 2019) that participants exert more cognitive effort when the reward rate is low, we demonstrate that recent performance accuracy can fully explain this effect. This study suggests that evidence previously used to support the idea that environmental (average) reward rate modulates cognitive effort should instead be interpreted as showing that accuracy rate plays that role. However, this does not undermine the theoretical suggestion that environmental reward rate acts as a heuristic for assessing the opportunity costs of cognitive resources. This still very intriguing idea should be explored in future studies investigating the hypothesized relationship between cognitive effort and environmental richness, while carefully ensuring the efficacy of reward rate manipulations.

## Open Practices Statement

This study was pre-registered on the Open Science Framework (OSF; https://doi.org/10.17605/OSF.IO/QET4M). All code and data are available at https://data.ru.nl/collections/di/dcc/DSC_2020.00081_234.

## Notes

### Competing Interest Statement

The authors have declared no competing interest.

### Funding

- Dutch Research Council 'Vidi' grant: NWO 016.175.450

