## SupplementalMaterials for "Dissecting the contribution of recent reward versus recent performance history on cognitive effort allocation"

### Supplemental Materials

#### Supplemental methods

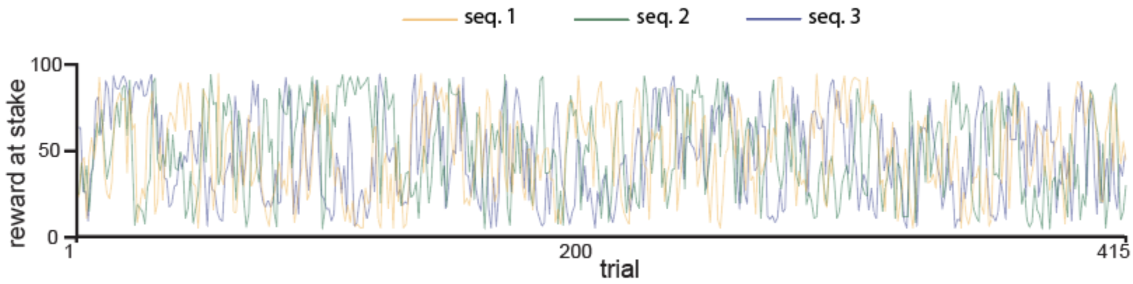

**Figure S1.** The three reward rate sequences used in the task, which were counterbalanced across participants.

#### Supplemental results

Figure S2 shows the basic effects of previous accuracy and previous congruency on accuracy and reaction times, as reported in the main text.

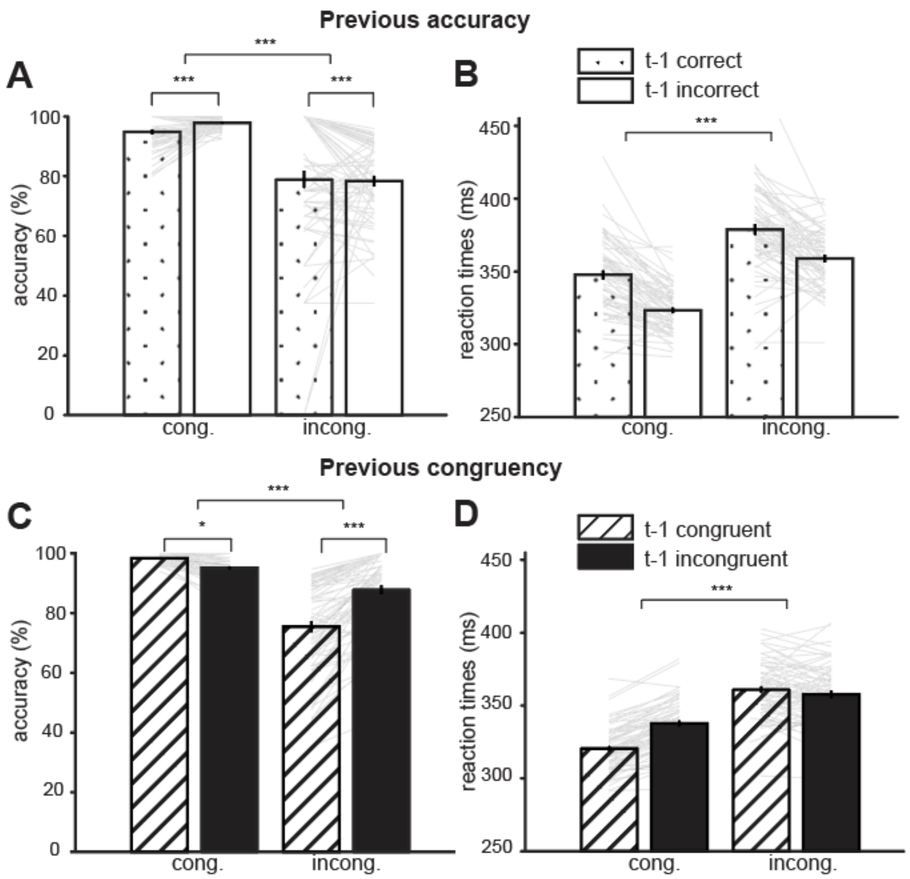

**Figure S2. Previous accuracy and previous congruency. A.** Accuracy and **B.** reaction times for congruent and incongruent trials, for previous correct (dots) and previous incorrect (white, no dots) trials. There was a significant effect of previous error on accuracy (β = -0.42, p = $1x10^{-14}$) and reaction times (β = 8.72, p = $1x10^{-12}$): participants were overall less accurate and slower after an incorrect response. **C.** Accuracy and **D.** reaction times for congruent and incongruent trials, for previous congruent (lines) and previous incongruent (black, no lines) trials. There was a significant effect of previous congruency on both accuracy (β = -0.26, p = $2x10^{-7}$) and reaction times (β =-4.69, p = $1x10^{-13}$): participants were overall less accurate and faster when the previous trial was congruent compared to when the previous trial was incongruent.

#### Absence of reward rate effect on RT

Following Lin et al. (2022), the learning rate was estimated on reaction times on the correct, congruent trials. To examine this model estimation, we performed a grid search analysis across 501 learning rates, performing the same regression analysis used to estimate the learning rate. Figure S3 shows the AIC minus the lowest AIC value for each model, demonstrating that the AIC remains approximately the same across the range of learning rates, further proving the absence of response time effects in the data.

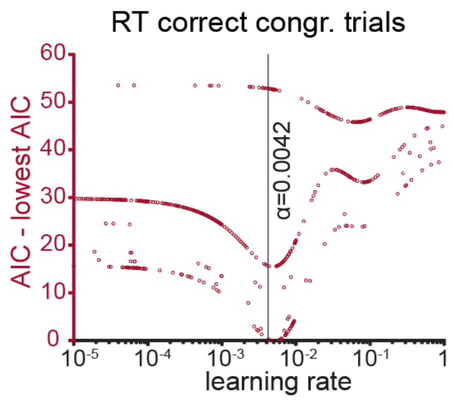

**Figure S3. Grid search analysis RT regression for correct, congruent trials.** These trials were also used for the learning rate estimation that resulted in an estimated learning rate of 0.0042. Learning rate was varied between 0.00001 and 1. AIC minus the lowest AIC value is shown in red. AIC remains approximately the same across the range of learning rates.

#### Accuracy analysis reward rate (RR) congruent trials

Table S1 shows the results from the mixed-effects regression accuracy analysis on the congruent trials, as reported in the main text.

| **Coefficient** | **β Estimate (SE)** | | **p-value** | |
|---|---|---|---|---|
| | **α1** | **α2** | **α1** | **α2** |
| (intercept) | 4.10 (0.12) | 4.11 (0.12) | $< 2x10^{-16}$ | $< 2x10^{-16}$ |
| Reward rate | -0.0048 (0.056) | -0.00026 (0.057) | 0.9 | 1.0 |
| Stake | 0.015 (0.052) | 0.018 (0.054) | 0.8 | 0.7 |
| RT | -0.69 (0.058) | -0.69 (0.059) | $< 2x10^{-16}$ | $< 2x10^{-16}$ |
| Congruency (t-1) | 0.47 (0.065) | 0.47 (0.065) | $9x10^{-13}$ | $1x10^{-14}$ |
| Error (t-1) | -0.17 (0.075) | -0.17 (0.075) | 0.026 | 0.026 |
| Response rep. | -0.11 (0.057) | -0.11 (0.058) | 0.064 | 0.062 |
| Trial number | -0.081 (0.058) | -0.078 (0.060) | 0.2 | 0.2 |

**Table S1. Accuracy - congruent trials**. Results from the mixed-effects logistic regression analysis examining effects on accuracy for the congruent trials. Results are consistent across two learning rates: α1 = 0.0031, α2 = 0.0042

#### Accuracy rate (AR) analysis

Here, we report an overview of the accuracy and reaction time regression results with accuracy rate, as reported in the main text.

| **Coefficient** | **β Estimate (SE)** | **p-value** |
|---|---|---|
| (intercept) | 2.51 (0.10) | $< 2x10^{-16}$ |
| Accuracy rate | -0.36 (0.04) | $< 2x10^{-16}$ |
| Stake | -0.011 (0.033) | 0.76 |
| RT | 0.47 (0.053) | $< 2x10^{-16}$ |
| Congruency | 1.47 (0.073) | $< 2x10^{-16}$ |
| Congruency (t-1) | -0.24 (0.052) | $3x10^{-6}$ |
| Error (t-1) | -0.46 (0.061) | $5x10^{-14}$ |
| Response rep. | -0.084 (0.032) | 0.009 |
| Trial number | 0.048 (0.047) | 0.3 |
| Accuracy rate x congruency | 0.062 (0.037) | 0.094 |
| RT x congruency | -1.27 (0.065) | $< 2x10^{-16}$ |

**Table S2. Accuracy**. Results from the mixed-effects logistic regression analysis examining effects on accuracy for all trials. Accuracy rate is calculated with learning rate 0.0031.

| Coefficient | β Estimate (SE) | p-value |
|---|---|---|
| (intercept) | 4.10 (0.12) | $< 2x10^{-16}$ |
| Accuracy rate | -0.25 (0.052) | $1x10^{-8}$ |
| Stake | 0.011 (0.054) | 0.8 |
| RT | -0.69 (0.052) | $< 2x10^{-16}$ |
| Congruency (t-1) | 0.48 (0.065) | $3x10^{-13}$ |
| Error (t-1) | -0.21 (0.077) | 0.007 |
| Response rep. | -0.11 (0.057) | 0.054 |
| Trial number | -0.064 (0.067) | 0.3 |

**Table S3. Accuracy – congruent trials**. Results from the mixed-effects logistic regression analysis examining effects on accuracy for the congruent trials. Accuracy rate is calculated with learning rate 0.0031.

| Coefficient | β Estimate (SE) | p-value |
|---|---|---|
| (intercept) | 2.11 (0.17) | $< 2x10^{-16}$ |
| Accuracy rate | -0.40 (0.045) | $< 2x10^{-16}$ |
| Stake | -0.0063 (0.043) | 0.9 |
| RT | 1.85 (0.11) | $< 2x10^{-16}$ |
| Congruency (t-1) | -0.84 (0.072) | $< 2x10^{-16}$ |
| Error (t-1) | -0.75 (0.11) | $2x10^{-12}$ |
| Response rep. | -0.10 (0.045) | 0.032 |
| Trial number | 0.15 (0.056) | 0.007 |

**Table S4. Accuracy – incongruent trials**. Results from the mixed-effects logistic regression analysis examining effects on accuracy for the incongruent trials. Accuracy rate is calculated with learning rate 0.0031.

| Coefficient | β Estimate (SE) | p-value |
|---|---|---|
| (intercept) | 352.37 (2.17) | $< 2x10^{-16}$ |
| Accuracy rate | -1.05 (0.56) | 0.07 |
| Stake | -0.38 (0.34) | 0.3 |
| Congruency | -17.69 (0.69) | $< 2x10^{-16}$ |
| Congruency (t-1) | -4.84 (0.50) | $5x10^{-14}$ |
| Error (t-1) | 8.33 (0.98) | $3x10^{-12}$ |
| Response rep. | 1.69 (0.69) | 0.017 |
| Trial number | 1.01 (0.66) | 0.1 |
| ITI | -5.60 (0.47) | $< 2x10^{-16}$ |

**Table S5. Reaction times**. Results from the mixed-effects linear regression analysis examining effects on reaction times for all correct trials. Accuracy rate is calculated with learning rate 0.0031.

| Coefficient | β Estimate (SE) | p-value |
|---|---|---|
| (intercept) | 2.14 (0.18) | $< 2x10^{-16}$ |
| Accuracy rate | -0.40 (0.050) | $9x10^{-11}$ |
| Reward rate residuals | -0.017 (0.050) | 0.8 |
| Stake | -0.012 (0.045) | 1.0 |
| RT | 1.87 (0.11) | $< 2x10^{-16}$ |
| Congruency t-1 | -0.85 (0.074) | $< 2x10^{-16}$ |
| Error t-1 | -0.74 (0.11) | $2x10^{-5}$ |
| Response rep. | -0.094 (0.045) | 0.1 |
| Trial number | 0.14 (0.057) | 0.047 |

**Table S6. Accuracy – incongruent trials, with accuracy rate and reward rate residuals.** Results from the mixed-effects logistic regression analysis examining effects on accuracy for incongruent trials, with both accuracy rate and the reward rate residuals included in the model. Accuracy rate and reward rate are calculated with learning rate 0.0031.

| Coefficient | β Estimate (SE) | p-value |
|---|---|---|
| (intercept) | 2.65 (0.14) | $< 2x10^{-16}$ |
| Accuracy rate | -2.28 (0.46) | $6x10^{-7}$ |
| RT | 0.79 (0.086) | $< 2x10^{-16}$ |
| Congruency | 1.55 (0.10) | $< 2x10^{-16}$ |
| Congruency (t-1) | -0.27 (0.095) | 0.004 |
| Error (t-1) | -0.50 (0.13) | 0.0001 |
| Response rep. | -0.011 (0.068) | 0.9 |
| Trial number | 2.19 (0.10) | $2x10^{-6}$ |
| Accuracy rate x congruency | -0.17 (0.10) | 0.091 |
| RT x congruency | -1.45 (0.087) | $< 2x10^{-16}$ |

**Table S7. Accuracy – preliminary trials**. Results from the mixed-effects logistic regression analysis examining effects on accuracy for all preliminary trials. Accuracy rate is calculated with learning rate 0.0031.

| Coefficient | β Estimate (SE) | p-value |
|---|---|---|
| (intercept) | -4.90 (0.24) | $< 2x10^{-16}$ |
| Accuracy rate | -2.52 (0.83) | 0.002 |
| RT | -0.51 (0.11) | $1x10^{-6}$ |
| Congruency (t-1) | 0.86 (0.14) | $4x10^{-10}$ |
| Error (t-1) | 0.30 (0.18) | 0.1 |
| Response rep. | -0.29 (0.11) | 0.009 |
| Trial number | 2.30 (0.83) | 0.006 |

**Table S8. Accuracy – congruent preliminary trials**. Results from the mixed-effects logistic regression analysis examining effects on accuracy for the congruent preliminary trials. Accuracy rate is calculated with learning rate 0.0031.

| Coefficient | β Estimate (SE) | p-value |
|---|---|---|
| (intercept) | 2.32 (0.26) | $< 2x10^{-16}$ |
| Accuracy rate | -2.38 (0.68) | 0.0004 |
| RT | 2.83 (0.18) | $< 2x10^{-16}$ |
| Congruency (t-1) | -1.51 (0.17) | $< 2x10^{-16}$ |
| Error (t-1) | -1.59 (0.22) | $4x10^{-13}$ |
| Response rep. | 0.17 (0.11) | 0.1 |
| Trial number | 2.45 (0.69) | 0.0003 |

**Table S9. Accuracy – incongruent preliminary trials**. Results from the mixed-effects logistic regression analysis examining effects on accuracy for the incongruent preliminary trials. Accuracy rate is calculated with learning rate 0.0031.

| Coefficient | β Estimate (SE) | p-value |
|---|---|---|
| (intercept) | 352.45 (2.89) | $< 2x10^{-16}$ |
| Accuracy rate | -18.75 (5.71) | 0.002 |
| Congruency | -18.26 (0.95) | $< 2x10^{-16}$ |
| Congruency (t-1) | -8.41 (0.89) | $9x10^{-14}$ |
| Error (t-1) | 20.99 (2.09) | $1x10^{-14}$ |
| Response rep. | -0.079 (1.12) | 0.9 |
| Trial number | 21.60 (5.66) | 0.0003 |
| ITI | -0.38 (0.68) | 0.6 |

**Table S10. Reaction times – preliminary trials**. Results from the mixed-effects linear regression analysis examining effects on reaction times for all correct preliminary trials. Accuracy rate is calculated with learning rate 0.0031.

#### Results replicate in pilot data set with 600ms response deadline

The Simon task used by Otto & Daw (2019) employed a 600 ms. response deadline, with 10% of trials having a response deadline of 500 ms. We first tested 33 participants with these same timings, but summary statistics showed that participants performed at ceiling level (accuracy congruent trials: 95.0% SD= 3.1, range: 90.4% - 99.5%, accuracy incongruent trials: 87.8% SD = 9.3, range: 63.0% - 99.1%). This high performance level compared to the results from Otto & Daw (2019) can likely be explained by participants performing the experiment in the lab where there were no distractions and participants were monitored by the experimenter, compared to doing the experiment online. To make performance equal to performance in the Otto & Daw (2019) paper, we shorted the response deadline to 500 ms. (with 10% of trials having a response deadline of 400 ms., see Methods).

We analysed the pilot data set to examine whether the results from the main dataset were present, despite the dataset comprising of fewer participants and participants performing at ceiling level. We analysed the data with the learning rate of 0.0031 (Otto & Daw, 2019). Similar to the main dataset, we find that participants are less accurate when the reward rate is high, and this effect almost reaches significance (β = -0.2, p = 0.052, Table S11). Unlike the main analysis, there was no interaction between reward rate and congruency (β = -0.009, p = 0.9), likely due to performance being high on both trial types. There was no effect of reward rate on response times (β = -0.1, p = 0.9, Table S12), nor were there effects of stake on accuracy (β = -0.007, p = 0.2) or response time (β = -1.0, p = 0.1). Like the main analysis, we find a highly significant effect of accuracy rate on accuracy (β = -0.5, p = $2x10^{-7}$, Table S13), but no effect of accuracy rate on reaction times (β = -0.3, p = 0.6, Table S14).

| Coefficient | β Estimate (SE) | p-value |
|---|---|---|
| (intercept) | 3.12 (0.20) | < $2x10^{-16}$ |
| Reward rate | -0.15 (0.070) | 0.052 |
| Stake | -0.0073 (0.067) | 0.2 |
| RT | 0.66 (0.11) | $6x10^{-8}$ |
| Congruency | 1.15 (0.15) | $1x10^{-9}$ |
| Congruency (t-1) | -0.17 (0.087) | 0.024 |
| Error (t-1) | -0.46 (0.17) | 0.007 |
| Response rep. | -0.072 (0.064) | 0.1 |
| Trial number | 0.051 (0.066) | 0.4 |
| Reward rate x congruency | -0.0090 (0.067) | 0.9 |
| RT x congruency | -1.45 (0.11) | < $2x10^{-16}$ |

**Table S11. Accuracy 600ms dataset**. Results from the mixed-effects logistic regression analysis examining effects on accuracy for all trials of the 600ms dataset. Reward rate is calculated with learning rate 0.0031.

| Coefficient | β Estimate (SE) | p-value |
|---|---|---|
| (intercept) | 392.53 (4.84) | < $2x10^{-16}$ |
| Reward rate | -0.14 (0.98) | 0.9 |
| Stake | -0.99 (0.63) | 0.1 |
| Congruency | -27.08 (1.49) | < $2x10^{-16}$ |
| Congruency (t-1) | -4.82 (0.66) | $9x10^{-10}$ |
| Error (t-1) | 9.46 (1.79) | $7x10^{-6}$ |
| Response rep. | 0.88 (0.94) | 0.2 |
| Trial number | -0.40 (1.25) | 0.3 |
| ITI | -6.56 (0.72) | $2x10^{-11}$ |

**Table S12. Reaction times 600ms dataset.** Results from the mixed-effects linear regression analysis examining effects on reaction times for all correct trials of the 600ms dataset. Reward rate is calculated with learning rate 0.0031.

| Coefficient | β Estimate (SE) | p-value |
|---|---|---|
| (intercept) | 3.12 (0.21) | $< 2x10^{-16}$ |
| Accuracy rate | -0.49 (0.094) | $2x10^{-7}$ |
| Stake | -0.021 (0.067) | 0.8 |
| RT | 0.66 (0.11) | $2x10^{-9}$ |
| Congruency | 1.16 (0.15) | $7x10^{-14}$ |
| Congruency (t-1) | -0.14 (0.086) | 0.099 |
| Error (t-1) | -0.55 (0.17) | 0.002 |
| Response rep. | -0.061 (0.064) | 0.3 |
| Trial number | -0.15 (0.087) | 0.082 |
| Accuracy rate x congruency | -0.012 (0.088) | 0.9 |
| RT x congruency | -1.47 (0.11) | $< 2x10^{-16}$ |

**Table S13. Accuracy 600ms dataset**. Results from the mixed-effects logistic regression analysis examining effects on accuracy for all trials of the 600ms dataset. Accuracy rate is calculated with learning rate 0.0031.

| Coefficient | β Estimate (SE) | p-value |
|---|---|---|
| (intercept) | 392.47 (4.87) | $< 2x10^{-16}$ |
| Accuracy rate | -0.32 (0.69) | 0.6 |
| Stake | -0.95 (0.63) | 0.1 |
| Congruency | -27.09 (1.50) | $< 2x10^{-16}$ |
| Congruency (t-1) | -4.83 (0.65) | $2x10^{-9}$ |
| Error (t-1) | 9.39 (1.79) | $1x10^{-5}$ |
| Response rep. | 0.86 (0.94) | 0.4 |
| Trial number | -0.86 (0.94) | 0.4 |
| ITI | -6.52 (0.72) | $1x10^{-10}$ |

**Table S14. Reaction times 600ms dataset.** Results from the mixed-effects linear regression analysis examining effects on reaction times for all correct trials of the 600ms dataset. Accuracy rate is calculated with learning rate 0.0031.

### Discussion

#### Learning rate estimation

We explored the robustness of the effects of reward / performance rate, and their dependence on fitting specific learning rates. For both reward and accuracy rates, we found that these explained cognitive effort at a range of low learning rates that reflected a long-range average reward/performance. This shows that the exact learning rate being used does not matter, as long as it is relatively low.

#### Considerations and future directions for successful reward rate manipulations

We suggest in the main manuscript that in order to study the effects of reward rate on performance, it is crucial to ensure that participants process the stake manipulation. Instead of focusing on the somewhat obfuscated reward at stake, participants may have directed their attention towards internal motivation, such as the desire to avoid mistakes (Hajcak & Foti, 2008). We believe that increasing the reward salience and reducing reward granularity could potentially address this. Regarding reward salience, a number of participants anecdotally reported that they were not paying attention to the stakes. Participants may have been driven by the guaranteed large monetary reward at the end of the task instead of the smaller, effort-contingent rewards during task performance (Grogan et al., 2020). Manipulations could include larger trial-by-trial rewards with a pay-out of only a few trials. Regarding stake granularity, in the studies not reporting an effect of reward at stake, including this study, stakes were varied at a high granularity, e.g. any integers between 1 and 100 (Beierholm et al., 2013; Devine et al., 2021; Guitart-Masip et al., 2012; Lin et al., 2022; Otto & Daw, 2019). These may be cognitively demanding to process (e.g. compare 38 vs 83, versus comparing 40 and 80 points). In contrast, when varying reward stakes between two levels in a similar conflict task did modulate responding, such that, responses were invigorated under high reward stakes (Hofmans et al., 2020).
